# Respiratory coupling between prefrontal cortex and hippocampus of rats anesthetized with urethane in theta and non-theta states

**DOI:** 10.1101/2021.03.31.437980

**Authors:** Rola Mofleh, Bernat Kocsis

## Abstract

Respiratory modulation of forebrain activity, long considered hard to reliably separate from breathing artifacts, has been firmly established in recent years using a variety of advanced techniques. Respiratory related oscillation (RRO) is derived from rhythmic nasal airflow in the olfactory bulb (OB) and is conveyed to higher order brain networks, including the prefrontal cortex (PFC) and hippocampus (HC), where it may potentially contribute to communication between these structures by synchronizing their activities at the respiratory rate. RRO was shown to change with sleep-wake states, it is strongest in quiet waking, somewhat less in active waking, characterized with theta activity in the HC, and absent in sleep. The goal of this study was to test RRO synchronization between PFC and HC under urethane anesthesia where theta and non-theta states spontaneously alternate. We found that in theta states, PFC-HC coherences significantly correlated with OB-HC but not with OB-PFC, even though RRO was stronger in PFC than in HC. In non-theta states, PFC-HC synchrony correlated with coherences connecting OB to either PFC or HC. Thus, similar to freely behaving rats, PFC-HC synchrony at RRO was primarily dependent on the response of HC to the common rhythmic drive, but only in theta state. The findings help outlining the value and the limits of applications in which urethane-anesthetized rats can be used for modeling the neural mechanisms of RRO in behaving animals.

## Introduction

Respiratory modulation of forebrain activity, long considered hard to reliably separate from breathing artifacts, has been firmly established in recent years ^1^, using a variety of advanced techniques, including single unit firing^2–4^, phase modulation of local gamma activity ^2, 3, 5–7^, and current source density analysis ^4, 8^. Many of these observations in awake or sleeping animals were preceded, supplemented, or verified under urethane anesthesia ^9, 10^, which besides providing a stable preparation, allows additional manipulations, e.g. tracheostomy, and lacks behavioral confounds related to e.g. sniffing. Importantly, different types of brain oscillations were shown to survive anesthesia with urethane better than with other anesthetics ^11–17^.

In freely moving animals, neural oscillations are related to and are intricately modified by behavior. Hippocampal (HC) theta rhythm for example is mainly associated with exploratory behavior and gross movements but may also occur in awake immobility, elicited by a range of sensory inputs which are hard to account for; in their classic paper for example, Kramis et al.^14^ reported theta activity in reaction even to “changes in the experimenter’s facial expression”. Since stable theta oscillations are present under urethane anesthesia however, this model has been widely used for mechanistic investigations of generating this rhythm ^18–27^ and related neuropharmacology ^28–33^.

Respiratory related oscillations (RRO) are generated in the olfactory bulb (OB), monitoring the nasal airflow, and are then conveyed to forebrain networks, including prefrontal cortex (PFC) and HC ^1^. Relatively stable slow (~2 Hz) RRO, present outside of the transient sniffing episodes, strongly depends on sleep-wake states, as shown in studies monitoring respiration along with polysomnography over all sleep-wake states in the same animal ^7, 34, 35^ as well as in studies focusing on selected states ^2, 8, 36^. It is strongest in waking, especially at rest associated with non-theta activity in the HC. Slow RRO also shows interregional differences; it is stronger in cortical leads, including PFC, in contrast to fast RRO, stronger in HC during sniffing ^8, 34, 37, 38^.

The frequencies of the relatively stable slow and the more transient fast RRO in rodents overlap, respectively, with delta and theta oscillations, intrinsically generated by brain networks. Rhythmic synchronizations of HC and PFC at these frequencies are thought to mediate key cognitive functions, and disruptions of HC-PFC coupling were implicated in psychiatric diseases ^39–42^. It was proposed that theta and delta oscillations may serve as parallel channels of communication between the HC and the PFC ^43^ and that RRO may contribute to this communication in a state-dependent manner ^34^. Regular ~2 Hz oscillations in the PFC also occur in rats anesthetized with urethane ^44^ where theta rhythm has also been routinely recorded in HC ^18^, spontaneously alternating with wide-band delta (large irregular activity ^14^). The goal of this study was to test RRO synchronization between PFC and HC in this preparation, in theta and non-theta states.

## Methods

Experiments were performed on 10 rats (Sprague-Dawley, male, 360–560g, Charles River Laboratories) under urethane anesthesia. All procedures were approved by the Institutional Animal Care and Use Committee of Beth Israel Deaconess Medical Center and carried out in accordance with relevant guidelines and regulations. The study and reporting also adheres to ARRIVE guidelines.

### Experimental procedures

Urethane (i/p; 1.2-1.5 g/kg of 65-80% solution) was given in two doses, an hour apart with ketamine supplemented (3.5 to 5 mg/kg) if necessary by periodically monitoring reaction to tail pinch thorough the experiment. The body temperature was maintained by isothermal pad. First, a 10 mm cut was made on the right side of the abdomen through which the diaphragm was palpated with the edge of the forceps to find the right position for implantation of the electrodes. Two multi-threaded electrode wires were implanted and fixed with surgical glue for recording of the diaphragmal (dia) EMG. The incision was closed by 5-6 sutures. The dia electrodes were tunneled under the skin up to the skull. The second part of the surgery was performed in a stereotaxic frame, where the recording electrodes were implanted according to the Atlas of Paxinos ^45^. Stainless steel screw above the parietal cortex (AP: -3.5 mm, Lat: 2.5 mm from bregma) was inserted on the left side to record the cortical EEG, and two screws were inserted in the nasal bone (~5.0 mm anterior to bregma) and above the cerebellum to act as reference and ground electrodes. Two single electrodes (stainless steel wires) were implanted on each side (AP: +3.2 mm, Lat: ±0.5 mm, DV: -5.1 mm) to record the electrical activity in the PFC, a single electrode on the left side for the OB (AP: +8.5 mm, Lat: 1.5 mm, DV: -1.6 mm), and a pair of twisted wires with 1 mm between their tips in the HC on the right side (AB: -3.7, Lat: 2.2 mm, DV: -3.5 mm). All the electrode wires and screws were fixed to the skull with dental acrylic. The electrodes were connected to an amplifier (A-M systems) for data recording, filtered between 0.3 and 100 Hz, whereas the dia EMG filters had both low- and high-pass filters set at 100 Hz to diminish unwanted noise including respiratory movement artifacts and electrical noise from the heart. Recordings started an hour after electrode implantation and lasted 2-3 hours over spontaneous alternation of theta and non-theta states. Local field potentials (LFP) were recorded and saved in DASYLab 7.0 Acquisition System at 1 kHz sampling rate. For analysis, multiple segments were selected from stable episodes of theta and non-theta states (Table 1). After the experiment, the rats were perfused with phosphate buffered saline (PBS) followed by 10% buffered formalin (15-20 min) for histological (Cresyl Violet staining) identification of electrode locations.

**Table 1:**
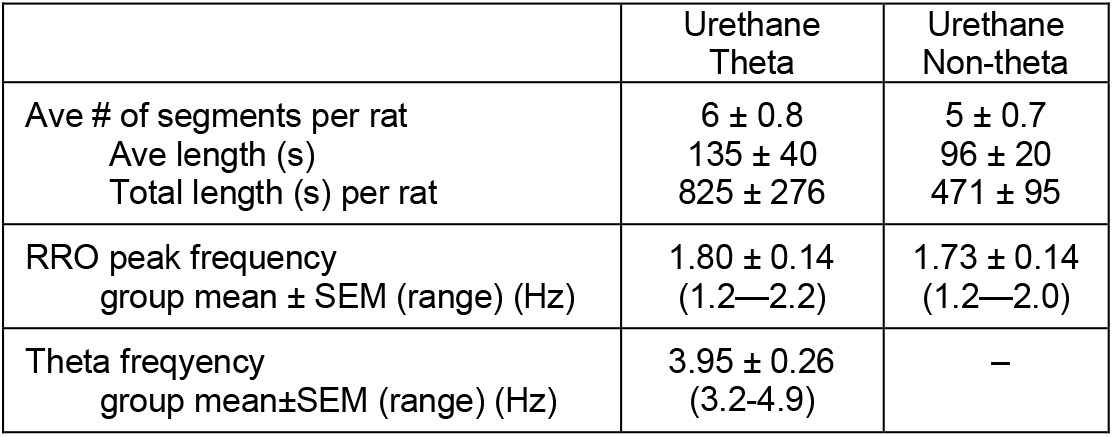
Length of segments and average RRO and theta frequency in different states

|  | Urethane Theta | Urethane Non-theta |
| --- | --- | --- |
| Ave # of segments per rat | 6 ± 0.8 | 5 ± 0.7 |
| Ave length (s) | 135 ± 40 | 96 ± 20 |
| Total length (s) per rat | 825 ± 276 | 471 ± 95 |
| RRO peak frequency<br>group mean ± SEM (range) (Hz) | 1.80 ± 0.14<br>(1.2–2.2) | 1.73 ± 0.14<br>(1.2–2.0) |
| Theta frequency<br>group mean ± SEM (range) (Hz) | 3.95 ± 0.26<br>(3.2–4.9) | – |

### Data analysis

was performed using Spike2 (Version 7- 06, Cambridge Electronic Devices) using procedures identical to those in a parallel study on freely moving rats ^34^. Dia EMG recordings were processed using built-in procedures of Spike2 to remove ECG contamination and to convert high-frequency EMG components in order to retrieve pure respiratory rhythm. To quantify neuronal synchronization between different structures we used pairwise coherences, calculated using a program (COHER.S2S) from the Spike2 library between four signal pairs, representing the potential transfer of the RRO signal to higher-order structures through the OB (i.e. dia with OB and OB with PFC and HC) and between these higher order structures (i.e. PFC with HC). Power spectra for dia EMG and HC were also calculated to identify the frequencies of spectral peaks of RRO and theta rhythm. Coherence values at RRO and theta frequencies calculated in multiple segments (Table 1) were averaged in each rat for theta and non-theta states and were then used in statistical analysis, including group averages and comparisons of peak coherences. Differences between coherences in different states were tested using Student's t-test after Fisher r to z transformation to obtain z-scored values with normal distribution. Correlation between pair-wise coherences were statistically tested using Excel’s T-DIST procedure using the formula p=TDIST(R*SQRT((N-2)/(1-R2)),N,1) where R is coherence and N is number of experiments, included.

## Results

Respiratory modulation of neural activity and of inter-regional coupling between OB, PFC, and HC networks was analyzed during theta and non-theta states, known to spontaneously alternate in rodents anesthetized with urethane. The length of these states varied from short episodes of 5-10 seconds to long continuous segments lasting 20-30 minutes. The switch between states was unpredictable (Fig. 1A); no tail pinch or any other intervention was used to trigger either state.

**Figure 1.**
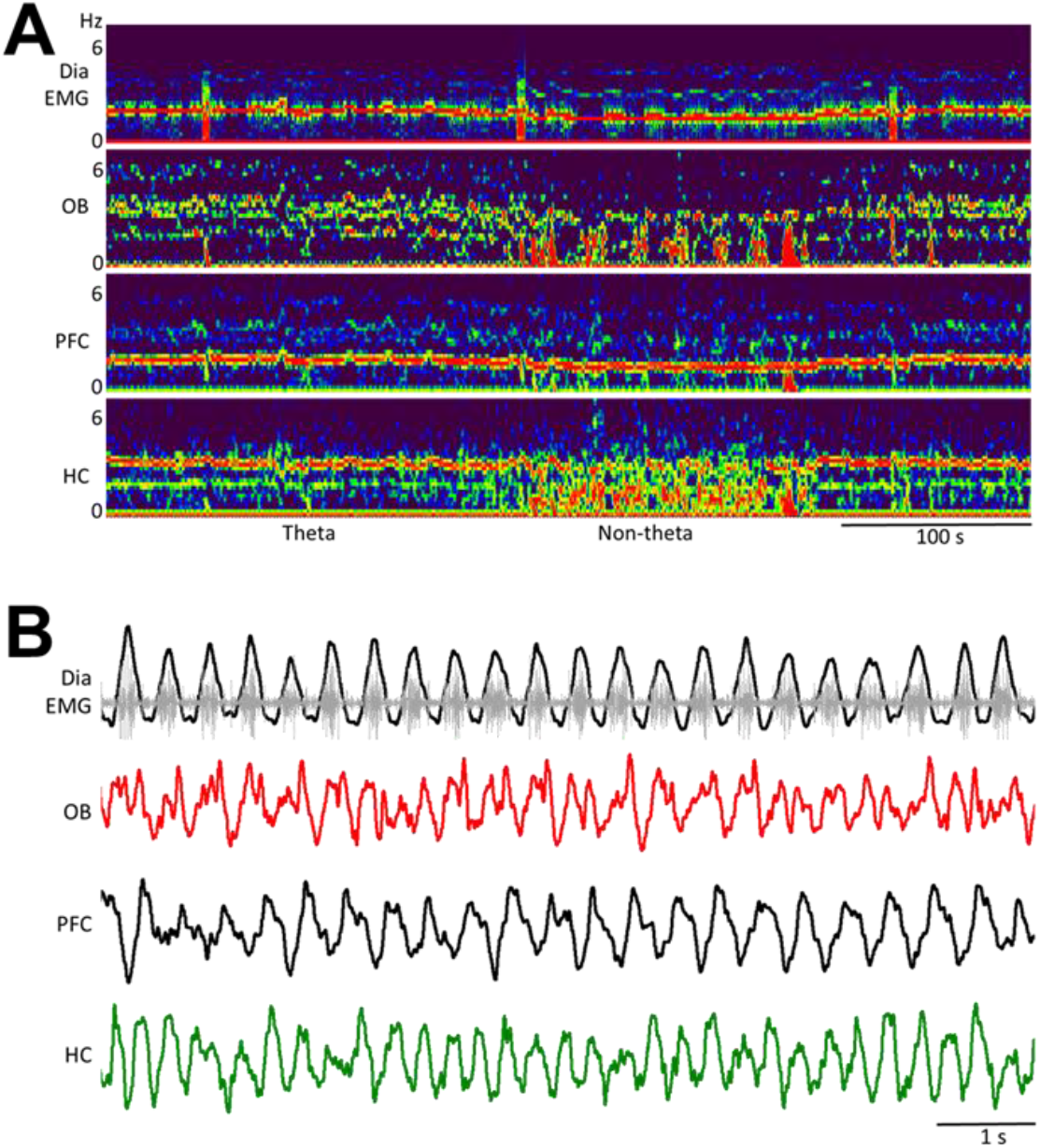
Sample recording of diaphragmal EMG along with LFPs in OB, PFC, and HC. **A**. time-frequency (0-7.5 Hz) plot showing alternation of theta - non-theta - theta states **B**. Traces of original recordings. Respiratory rhythm (black) was derived from diaphragmal EMG (grey)

To facilitate comparison, the investigation followed the structure used in a parallel study on freely behaving animals ^34^ based on analysis of multiple 80-150 s-long segments in different states. Thus, in each rat, multiple segments of this length scattered over the 2-3-hour recording were selected in theta and non-theta states (Table 1), and subjected to analysis of coherences between different structures and their correlations, at the respiratory frequency. To assess RRO synchronization across regions, pairwise coherences representing RRO transfer from rhythmic nasal airflow to the OB and then from OB to PFC and HC were compared in different states.

The animals were breathing spontaneously. The respiratory rate was below 2 Hz (Fig. 1), similar to that in unanesthetized rats during sleep ^7, 34, 35^, and remained stable over alternating theta and non-theta states (1.80±0.14 and 1.73±0.14 Hz, respectively; Table 1), resulting in a single sharp peak in the diaphragmal (dia) autospectra (Fig. 2A). Notably, RRO frequency was below that in waking (R1), even in “active states” under urethane. LFP in the OB correlated with this signal giving rise to RRO peaks in the dia-OB coherence spectra, also restricted to this narrow frequency range (Fig 2A).

**Figure 2.**
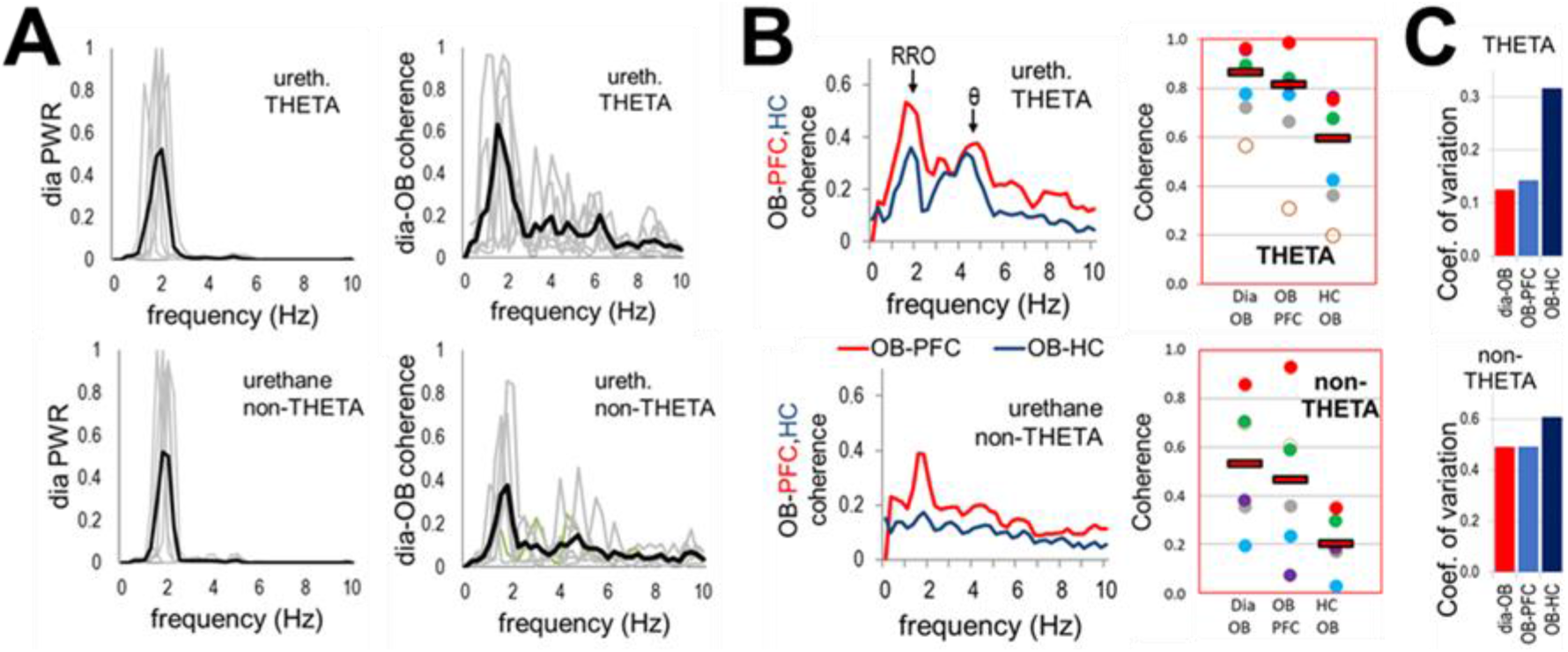
Comparison of state-dependent RRO coherences in PFC and HC transferred through OB in urethane-anesthetized rats. **A.** diaphragmal (dia) power and dia-OB coherence during theta and non-theta states. Dia-PWR, dia-OB coherence spectra are shown in individual experiments (grey) and averaged over the group (dark). **B.** Coherence between OB and higher order networks of PFC and HC. Average coherence spectra over RRO and theta frequencies (1-10 Hz; *left*) and peak coherence at RRO frequency (*right*; squares: group averages, dots: individual experiments; individual rats are marked using the same color in the two states). Note strong state dependence with nearly identical dia-OB and OB-PFC coherences in both states and considerably lower OB-HC coherence. **C.** Variability of coherence values in individual experiments. Note high coefficient of variation of OB-HC in theta state, twice exceeding dia-OB and OB-PFC.

RRO coupling appeared stronger in theta than in non-theta state (Fig. 2B), as shown by higher coherences between OB and all other signals (Table 2; p=0.012, 0.002, 0.007 for OB-dia, OB-PFC, OB-HC, respectively). RRO transmission through OB to forebrain structures also showed regional differences (Table 2). OB-HC coherence was relatively low, significantly lower than dia-OB coherence (p=0.01 and p=0.03, in the two states), whereas RRO was faithfully transmitted through the OB toward the PFC (p=0.37 and p=0.49). Examination of pairwise coherences in individual experiments further supported the pattern revealed by group averages; OB-HC coherence was lower than dia-OB and OB-PFC in all experiments, with no exception (see colored of dots in Fig. 2B). Coherence values varied between experiments in a wide range in all signal pairs, in both states. In theta state however, their deviations relative to the mean (coefficient of variation; CV) was higher for OB-HC (CV=0.32) than for dia-OB and OB-PFC (CV=0.13 and 0.14, respectively), i.e. the variance stayed even (p>0.14; F-test) and did not follow significant differences in their corresponding group averages (Fig. 2C).

**Table 2.** Coherences connecting OB to dia and LFP signals of PFC and HC, at RRO frequency (*left*) and correlations (R^2^) between C(HC-PFC) and coherences connecting OB to other signals (*right*). Yellow background highlights significant correlations (p < 0.05) between pairwise coherences. C(X-Y): coherence between X and Y at the frequency of respiration.

| RRO Coherence | Urethane Theta | Urethane Non-theta |
| --- | --- | --- |
| C(Dia-OB) | 0.87 ± 0.06 | 0.53 ± 0.11 |
| C(OB-PFC) | 0.82 ± 0.09 | 0.47 ± 0.13 |
| C(OB-HC) | 0.60 ± 0.08 | 0.20 ± 0.05 |
| C(PFC-HC) | 0.70 ± 0.05 | 0.30 ± 0.08 |
| **Correlation ($R^2$) C(PFC-HC) with...** |  |  |
| ... C(Dia-OB) | 0.63 | 0.87 |
| ... C(OB-PFC) | 0.30 | 0.92 |
| ... C(OB-HC) | 0.72 | 0.88 |

Analysis of the variation between experiments revealed further that transmission through OB played a dominant role in maintaining the level of RRO in both structures. dia-OB coherences quantifying RRO input to the OB positively correlated with RRO coherences in the pathways connecting OB to the PFC and to the HC (R>0, p<0.05, R^2^ between 0.80-0.99; Fig. 3), indicating that the more RRO is derived by OB from rhythmic nasal airflow, the more it is transferred further, to downstream targets of the OB.

**Figure 3.**
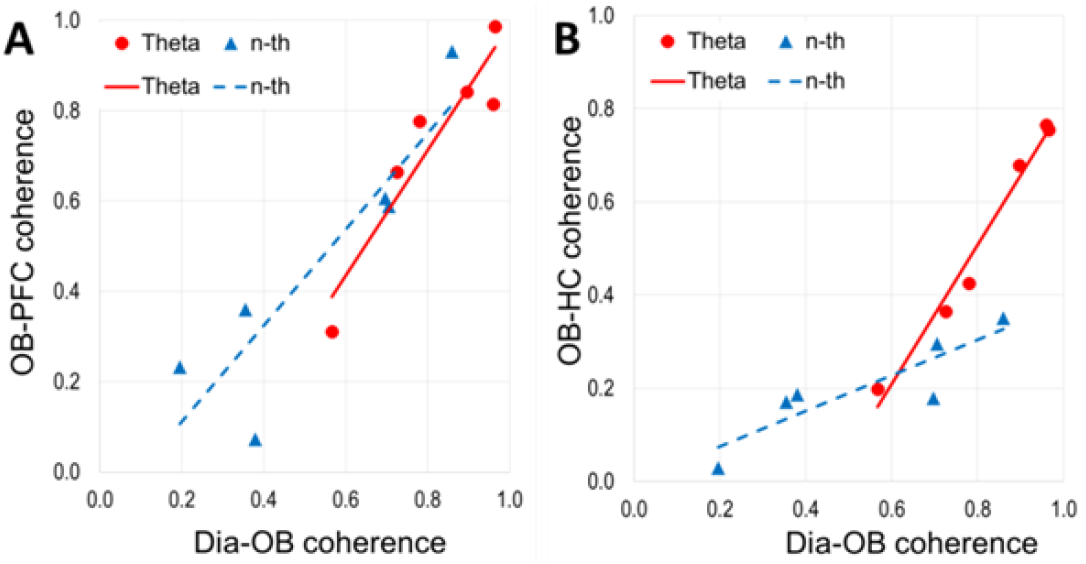
The relationship between peak RRO coherences connecting dia to OB and those connecting OB to PFC (A) and HC networks (B). Note significant (p<0.01) positive correlation of dia-OB with OB-PFC (R^2^=0.87 in theta and R^2^=0.80 in non-theta state) and OB-HC (R^2^=0.99 in theta and R^2^=0.78 in non-theta state).

RRO coupling was prominent between PFC and HC and was also found state dependent. It was consistently weaker in non-theta (0.30±0.08; Table 2) than theta state (0.70±0.05) where it was as high as the dominant theta coherence between the two structures (Fig. 4A). In non-theta state, individual variations in PFC-HC coherence strongly correlated with RRO conveyed from dia through OB to both PFC and HC (Fig. 4C) whereas in theta state RRO transmission to HC appeared dominant, strongly correlating with PFC-HC coherence (R^2^=0.72, p=0.02) compared with that to PFC (R^2^=0.30, p=0.16, Fig. 4B, Table 2).

**Figure 4.**
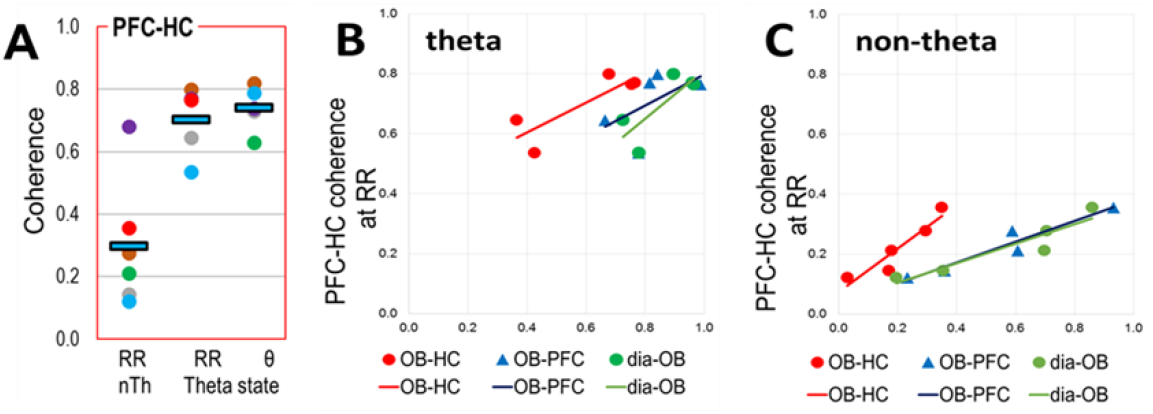
**A.** PFC-HC coherences under urethane anesthesia at RRO in non-theta and theta states and at θ frequency in theta states. Squares: group averages, dots: individual experiments (individual rats marked by the same colors as in Fig. 2.) **B-C.** Correlation between peak RRO coherences connecting PFC and HC vs. those between OB and dia, PFC, HC signals in theta and non-theta states. All correlations with PFC-HC were significant (<0.04 in theta and p<0.001) in non-theta state), except OB-PFC in theta state (R^2^=0.30, p=0.16).

## 4. Discussion

This study confirmed stable RRO in urethane-anesthetized rats ^10, 46^ and then proceeded to address the mechanism of RRO synchronization between higher order brain structures using inter-regional coherences and their correlations. First, we found significant correlation between coherences connecting OB activity to respiration on one hand and to PFC and HC activities on the other, in both theta and non-theta states. Second, in theta states, PFC-HC coherences significantly correlated with OB-HC but not with OB-PFC, even though RRO was stronger in PFC than in HC. In non-theta states, PFC-HC synchrony correlated with coherences connecting OB to either PFC or HC. Thus, similar to freely behaving rats ^34^, PFC-HC synchrony at RRO was primarily dependent on the response of HC to the common rhythmic drive, but only in theta state.

The major strength of this preparation, i.e. urethane-anesthetized rats, is that behavioral confounds of waking (e.g. locomotion, cognition) on one hand and fragmentation of sleep patterns (e.g. short REM sleep, never lasting longer than a few minutes in rodents) on the other are effectively eliminated. These advantages remain even in the face of substantial progress in recording technology and goes beyond the “convenience” provided by an immobile, stable preparation. It comes, however, with the price of uncertainties in the interpretation of alternating theta and non-theta states, weather they are analogous to differences between quiet vs. active waking or between slow wave sleep vs. REM sleep. Such concern is especially relevant for RRO which in freely behaving animals does not have clear association with theta; theta and RRO can occur either together (in active waking) or separately (theta in REM sleep and RRO in quiet waking).

Under urethane, RRO was present in both theta and non-theta states. RRO coupling was stronger in urethane-theta compared with non-theta state which however cannot be considered as evidence for a functional link between the two oscillations. In fact, their co-occurrence appeared opposite to that in awake animals where RRO is weaker in theta states than in quiet waking. RRO and theta rhythm were also shown mechanistically distinct and independent, even in conditions when their frequencies overlap. Besides their coherence with respiration and dependence on rhythmic nasal airflow ^1^, the two oscillations have different laminar profiles, they selectively entrain different gamma bands, and differ for example in sensitivity to cholinergic transmission ^3, 4, 10, 46^. Theta and non-theta states, frequently called “active”, and “passive” states, are associated with characteristic patterns of brainstem and cortical activity and related pharmacology ^18, 30, 47^, and therefore monitoring HC theta rhythm may be valuable in studies of RRO, but only as a good indicator of urethane-states.

Two further observations of this study help outlining the value and the limits of applications in which urethane-anesthetized rats can be used for modeling the neural mechanisms of RRO in behaving animals. The first concerns how respiratory rhythm is transmitted from mechanical sensory input in the OB to different forebrains structures, the second, how these structures respond to the OB input and how dynamic RRO coupling is established between them.

Under urethane, there is a faithful transfer of the respiratory rhythm through OB to both PFC and HC in both states, i.e. the more RRO is derived by OB from rhythmic nasal airflow, the more is transferred further, to PFC and HC. This matches the significant correlation between dia-OB and OB-PFC in active and quiet waking. However, strong RRO transmission through OB (i.e. dia-to-OB-to-HC) under urethane may be less valuable, as their relationship in freely moving rats is weaker, i.e. only amounts to a non-significant tendency ^34^. Furthermore, the results of this study indicate that modeling RRO coupling between higher-order forebrain structures, reported in freely behaving animals ^34^, can only use theta states under urethane anesthesia. Stable OB-PFC coherence contrasting low but highly variable OB-HC coherence in waking is only replicated in theta states under urethane. In this state, PFC-HC coupling significantly correlated with the OB-HC but not with OB-PFC coherence, indicating that HC access to the PFC output is dynamically regulated by the responsiveness of HC to the common rhythmic drive.

RRO mechanisms were widely explored earlier under urethane anesthesia ^10, 36, 46^ interpreting the results mostly in terms of modeling active and passive sleep states. Indeed, stable respiratory rate, along with several other parameters (e.g. genioglossus muscle atonia ^48^) in theta state resembles REM sleep more than AW. The results of this study however suggest a cautious approach when interpreting the results of RRO under urethane anesthesia. In anesthetized rats, RRO coherences in the dia-OB-PFC pathway in theta state appeared closer to those in QW than in the natural theta-dominated states of AW and REM sleep. In non-theta state they were higher than in SWS, the presumed unanesthetized analog of this state, and were instead in the range of chronic wake states. Between OB and HC, RRO coherence under urethane was higher in theta state than in any state of chronic experiments and low in non-theta states, in the range of chronic recordings. Lack of RRO under urethane was occasionally reported in previous studies, but only in deep anesthesia characterized by slow neocortical up-down transitions ^36, 46^, i.e. a pattern similar to natural slow wave sleep, where however RRO remains absent after switching to theta-rich REM sleep ^7, 34, 35^. Under urethane anesthesia, both OB-PFC and OB-HC coherences showed strong correlations with dia-OB coherence in both theta and non-theta states, again violating the correspondence, commonly expected on the basis of EEG signals, between urethane-theta and AW-REM on one hand and between urethane non-theta and QW-SWS on the other. These differences should be carefully considered in mechanistic studies of RRO mechanisms in the urethane model, in the future.

## Notes

### Competing Interest Statement

The authors have declared no competing interest.

